# Bright New Resources for Syphilis Research: Genetically Encoded Fluorescent Tags for *Treponema pallidum* and Sf1Ep Cells

**DOI:** 10.1101/2024.05.29.596454

**Authors:** Linda Grillová, Emily Romeis, Nicole A. P. Lieberman, Lauren C. Tantalo, Linda H. Xu, Barbara Molini, Aldo T. Trejos, George Lacey, David Goulding, Nicholas R. Thomson, Alexander L. Greninger, Lorenzo Giacani

**Affiliations:** Parasites and Microbes Programme, Wellcome Sanger Institute, Hinxton, UK; Department of Medicine, Division of Allergy and Infectious Diseases, University of Washington, Seattle (WA), USA; Department of Laboratory Medicine and Pathology, University of Washington, Seattle (WA), USA; Department of Global Health, University of Washington, Seattle (WA), USA; Faculty of Infectious and Tropical Diseases, London School of Hygiene & Tropical Medicine, London, UK; Vaccine and Infectious Disease Division, Fred Hutchinson Cancer Center, Seattle (WA), USA

**Keywords:** GFP-expressing *Treponema pallidum* subsp. *pallidum*, mCherry and BFP-expressing Sf1Ep cells, genetic manipulation, fluorescence microscopy

## Abstract

The recently discovered methodologies to cultivate and genetically manipulate *Treponema pallidum* subsp. *pallidum* (*T. pallidum*) have significantly helped syphilis research, allowing the *in vitro* evaluation of antibiotic efficacy, performance of controlled studies to assess differential treponemal gene expression, and generation of loss-of-function mutants to evaluate the contribution of specific genetic loci to *T. pallidum* virulence. Building on this progress, we engineered the *T. pallidum* SS14 strain to express a red-shifted Green Fluorescent Protein (GFP) and Sf1Ep cells to express mCherry and blue fluorescent protein (BFP) for enhanced visualization. These new resources improve microscopy- and cell sorting-based applications for *T. pallidum*, better capturing the physical interaction between the host and pathogen, among other possibilities. Continued efforts to develop and share new tools and resources are required to help our overall knowledge of *T. pallidum* biology and syphilis pathogenesis reach that of other bacterial pathogens, including spirochetes.

**Graphical abstract:** By employing genetic engineering, *T. pallidum* was modified to express GFP, and Sf1Ep cells to express mCherry on the cytoplasmic membrane and BFP in the nucleus. These new resources for syphilis research will facilitate experimental designs to better define the complex interplay between *T. pallidum* and the host during infection.

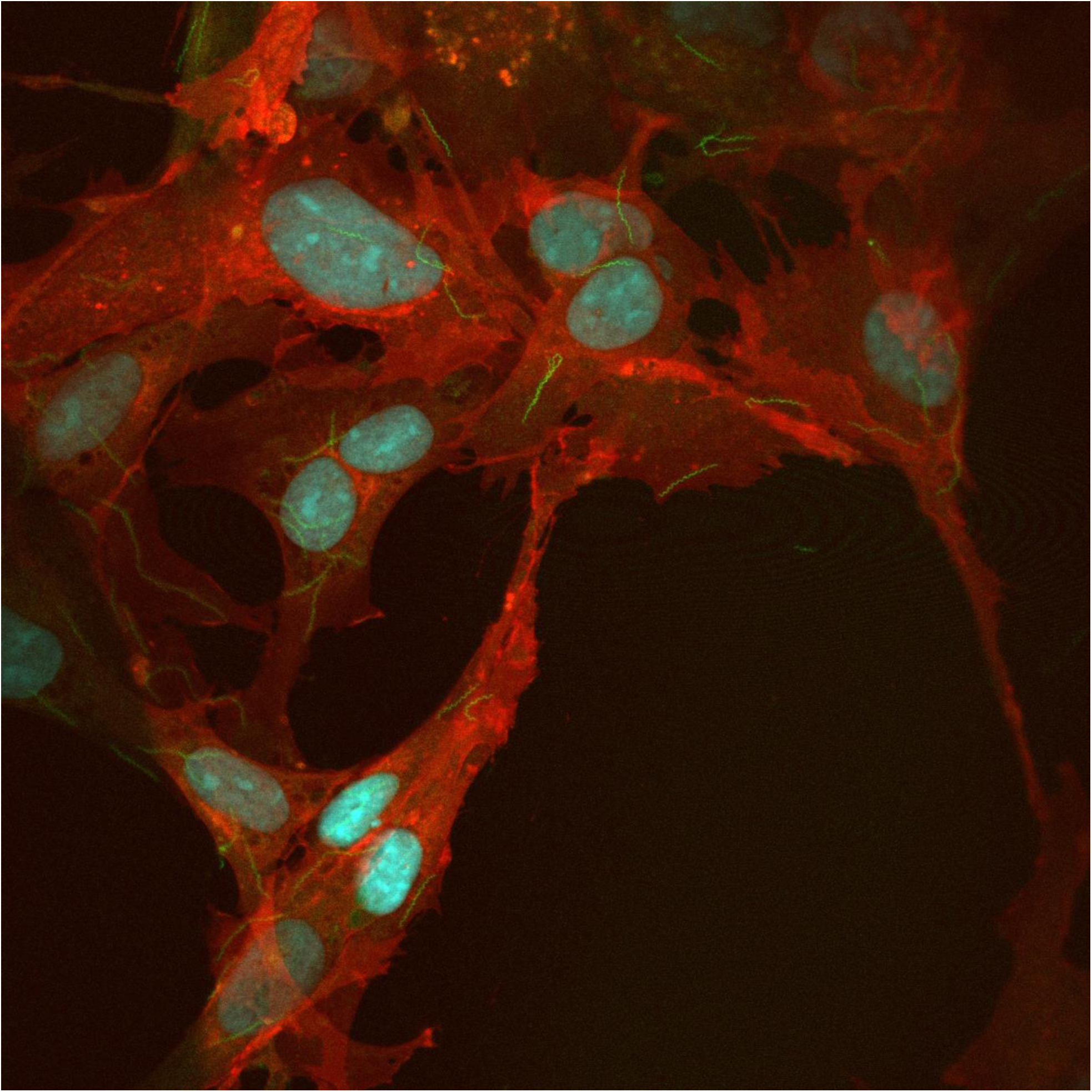

## INTRODUCTION

Basic syphilis research has faced significant obstacles in the past due to the inability to culture *Treponema pallidum* subsp. *pallidum* (*T. pallidum*) *in vitro*, which significantly hindered investigators’ ability to understand syphilis pathogenesis and *T. pallidum* biology. Since *T. pallidum* isolation in 1912 (1), researchers had to rely on the rabbit model of infection for a steady supply of viable *T. pallidum* cells for experiments and, even in laboratories willing to support this animal model, experimental procedures and designs were inevitably affected by the limited viability of the pathogen outside of a susceptible host. Two recent discoveries, however, marked a significant turning point in syphilis research. The first was defining the conditions for long-term *in vitro* propagation of *T. pallidum*, achieved through co-culture with rabbit skin epithelial Sf1Ep cells (2), and the second was the demonstration that *T. pallidum* could uptake exogenous DNA through exposure to a calcium chloride-based buffer and, therefore, be genetically manipulated (3).

The availability of an *in vitro* cultivation system has allowed, for example, the assessment of the efficacy of approved and new antibiotics (4–7), highly controlled studies to evaluate differences in gene expression in treponemes cultivated in parallel *in vitro* and *in vivo* (8), further characterization the *T. pallidum* hypervariable TprK virulence factor (9), and the development of an approach for genetic engineering of the syphilis agent (10).

Developing and sharing additional molecular tools for syphilis research remains a high priority, as they hold the key to improving our relatively limited understanding of the complex interplay between *T. pallidum* and the host during infection. Tools that facilitate pathogen visualization and detection are particularly useful. Here, we report two new resources for syphilis research, namely a genetically modified *T. pallidum* SS14 strain where the *tprA* (*tp0009*) pseudogene was replaced by a cassette for cytoplasmic expression of a red-shifted Green Fluorescent Protein (GFP)-Kan^R^ fusion, and Sf1Ep cells engineered to express mCherry in their cytoplasmic membrane and Blue Fluorescent Protein (BFP) in the nucleus. These novel tools will facilitate various research applications and projects ranging from fluorescence microscopy, cell sorting, and *in vivo* imaging following animal inoculation to study pathogen dissemination and transmission.

## RESULTS

### *T. pallidum* transformation, selection, and qualitative PCR

The pUC57-based plasmid used here contained the *gfp-kan*^R^ chimeric transgene preceded by the *tp0574* promoter. The transgene sequence was bounded by two homology arms corresponding to regions immediately upstream and downstream of the *tprA* (*tp0009*) pseudogene sequence (see methods). Wild-type *T. pallidum* SS14 (WT-SS14) treponemes were transformed by the introduction of this recombinant vector (File S1). *T. pallidum* transformants were selected in kanamycin-supplemented media. The WT-SS14 strain was also propagated in parallel, with and without kanamycin, as control. By the third passage, no WT-SS14 cells propagated in kanamycin-containing media were visible by darkfield microscopy (DFM). By passage #4, GFP-SS14 treponemes could be counted (1.3x10^8^/ml) in the well containing the transformed strain and kanamycin. Two weeks later, at passage #5, the culture was transferred from one well of a 24-well plate to six wells of a 6-well plate to be expanded. By passage #7 the yield of GFP-SS14 treponemes was comparable to the WT-SS14. Sub-culturing continued at weekly intervals with no interruption up to passage #31, when the strain was frozen after preliminary experiments were concluded. To confirm the replacement of *tprA* with the transgene sequence, a series of qualitative amplifications were performed (primers in Table 1) using genomic DNA extracted from GFP-SS14 cells. Firstly, using primers annealing to the *kan*^R^ and at sites within the *tprA* pseudogene sequence, no *tprA*-specific amplicon was observed using GFP-SS14 DNA that, however yielded a positive amplification for the *kan*^R^ target. DNA from the WT-SS14 strain showed the opposite amplification profile (Fig.1A). Amplification of the *tp0574* ORF was seen in both GFP-SS14 and WT-SS14 strain DNA (Fig.1A). Finally, primers targeting the regions upstream and downstream of the 5’ and 3’ *tprA* homology sequences also yielded an amplicon using both strains’ template DNA. The modest difference in predicted amplicon size for the WT *tprA* and the replacement *gfp-kan*^R^ chimeric transgene (174 bp) is difficult to fully appreciate following gel electrophoresis, due to the size and similarity in predicted amplicon sizes (4,469 bp for the GFP-SS14 vs 4,643 bp for the WT-SS14 strain) (Fig.1B). Hence, the amplified products were digested with XhoI to confirm the replacement of *tprA* locus with the transgene: the WT *tprA* gene does not contain a XhoI restriction site but the *kan*^R^ cassette within the transgene does (Fig.1B). Fluorescence of WT-SS14 and GFP-SS14 pellets measured at the time of harvest from *in vitro* harvested cells using a fluorescence reader demonstrated significantly higher signal from GFP-expressing treponemes compared to WT cells (Fig.1C).

**Figure 1:**
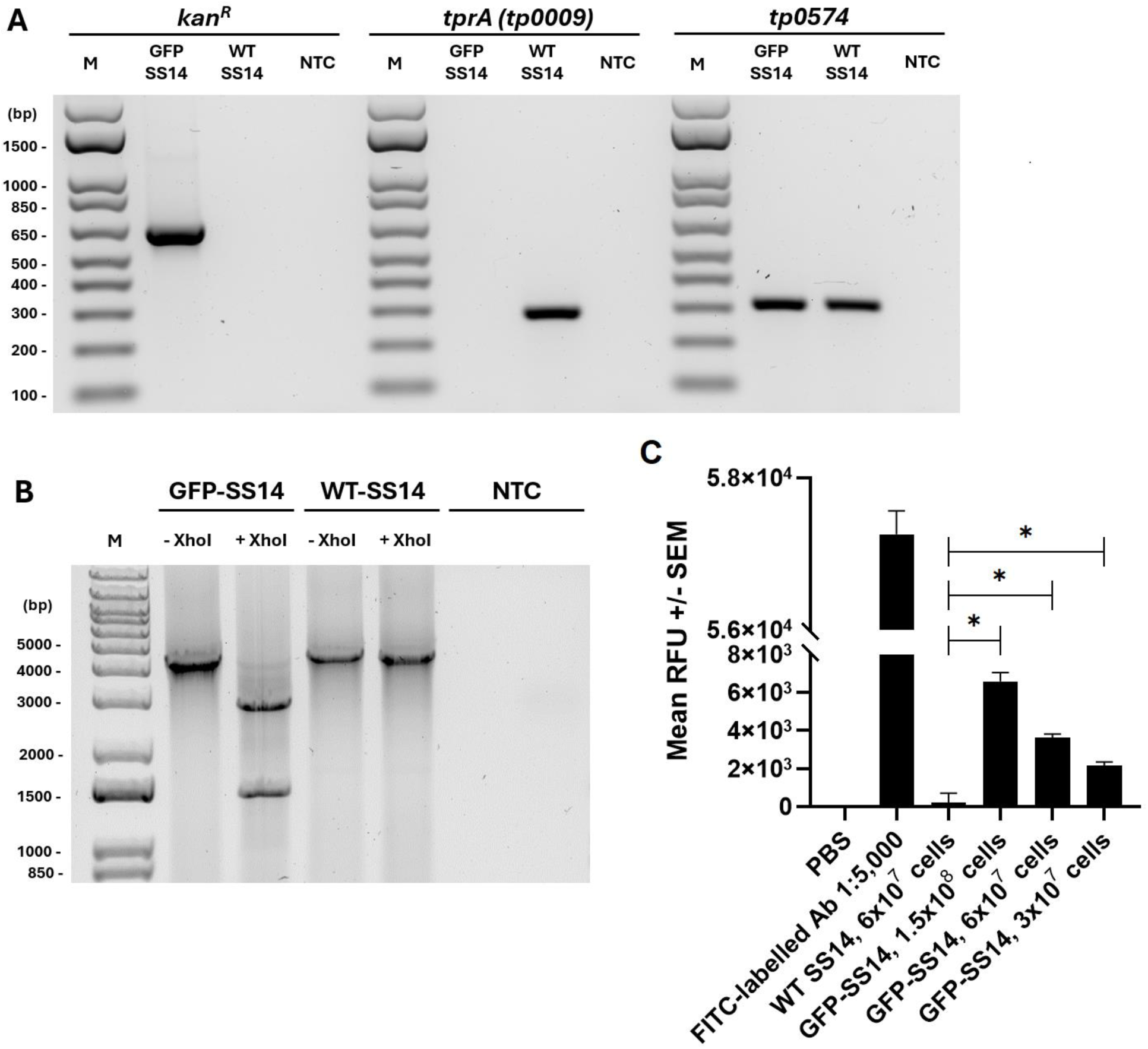
Qualitative PCR comparing the GFP-SS14 and the WT-SS14 strains. Amplification reactions using the primer combinations in Table 1 on DNA template from the GFP-SS14 and WT-SS14 strain harvested at passage #9 post-transformation. **(A)** Amplification using *kan*^R^ F/R primers, the *tprA* F/R primers, and the *tp0574* F/R primers. **(B)** XhoI digestion of amplicons generated using the left/right *tprA* flanking region F/R primers. M: molecular size marker (bp); NTC: no-template control. **(C)** Fluorescence measurement of WT-SS14 and GFP-SS14 cell pellets resuspended in PBS. RFU: Relative Fluorescence Units.

**Table 1.**
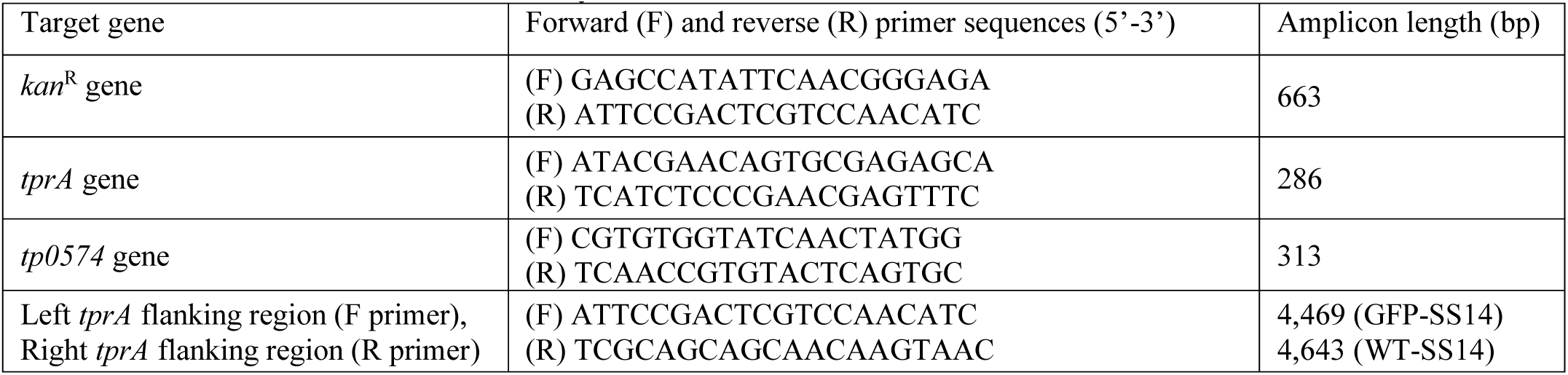
Primers used in this study.

### Whole-genome sequencing confirmation of GFP replacement of *tprA*

Whole-genome sequencing (WGS) was performed using a custom hybridization capture panel to enrich *T. pallidum* DNA to demonstrate the replacement of the *tprA* (*tp0009*) pseudogene by the *gfp-kan*^R^ chimeric transgene controlled by the *tp0574* promoter previously described by Weigel *et al.* (11). Results showed the absence of *tprA* in GFP-SS14 when read assembly was performed using the wild-type SS14 reference genome (CP004011.1; Fig.2A). However, when the assembly was guided by a template where *tprA* was replaced with the *tp0574* promoter/*gfp-kan*^R^ transgene, results showed replacement of *tprA* with the expected sequence (Fig.2B). Wild-type SS14 (WT-SS14), propagated in parallel to the GFP-SS14 strain, and sequenced as control, still carried *tprA* but, as expected, showed no reads mapping to the transgene (Fig.2C/D). Because our sequencing approach used enrichment probes based on wild-type *T. pallidum* genomes and did not include probes for the *kan*^R^ or *gfp* genes, coverage of the transgene was lower than targeted regions of the *T. pallidum* genome (Fig.2B), as seen previously (3, 10). These results confirmed the evidence previously attained by qualitative amplifications of the *tprA* locus and demonstrated the lack of WT-SS14 in the GFP-SS14 culture. An additional search for reads matching the plasmid backbone yielded no results, demonstrating lack of residual plasmid sequences in the culture. Further analyses failed to support that the transgene integrated anywhere else in the genome aside from the *tprA* locus. Strain sequences were deposited in GenBank with accession number CP148145 (GFP-SS14) and CP148144 (WT-SS14).

**Figure 2:**
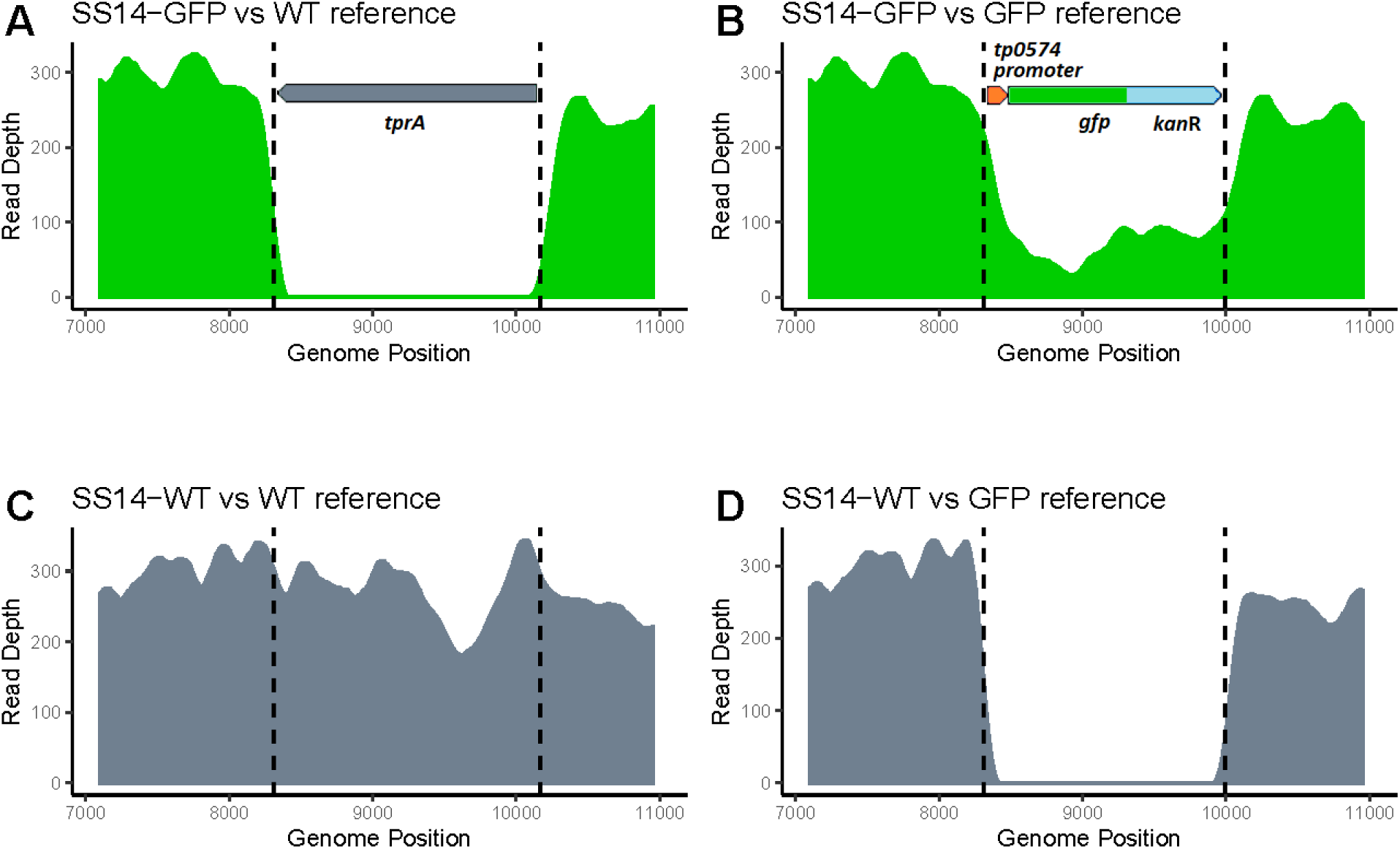
Whole genome sequencing of the GFP-SS14 strain. Genome sequencing confirms replacement of the *tprA* pseudogene with the *gfp-kan^R^* cassette. Assembled reads from GFP-SS14 **(A, B)** and the WT-SS14 **(C, D)** strains were mapped against the SS14 wild type reference **(A, C)** or the GFP-transgenic reference sequence **(B, D)**. The gap in coverage when SS14-GFP is mapped to the WT reference **(A)** demonstrates that no residual *tprA*-containing WT treponemes are present. Lower coverage **(B)** for the transgene, compared to the deleted sequence **(C)** is expected, and due to the lack of enrichment probes specific to the sequence replacing the *tprA* pseudogene.

### Fluorescence microscopy of GFP-expressing and wild-type *T. pallidum* on Sf1Ep cells

We demonstrated that expression of a single copy of a chromosomally integrated *gfp* gene is sufficient to detect bacteria by fluorescent microscopy. To facilitate the visualization, we have imaged the GFP-SS14 strain attached to the Sf1Ep-mCherry-BFP transduced cells (Fig.3, top panels). Recombinant lentivirus was used to deliver fluorescent markers into the genome of the Sf1Ep cells to permanently express mCherry in the membrane and BFP in nuclei (see Experimental Procedures). These modifications do not influence the ability of the Sf1Ep cells to support the growth of *T. pallidum* cells. As a control, wild-type *T. pallidum* cells were imaged alongside the Sf1Ep-mCherry-BFP transduced cells, which, as anticipated, showed no green fluorescence (Fig.3, middle panels), unless immunostaining of *T. pallidum* cells with a rabbit polyclonal antibody conjugated to FITC was performed following sample processing (Fig.3, bottom panels).

**Figure 3:**
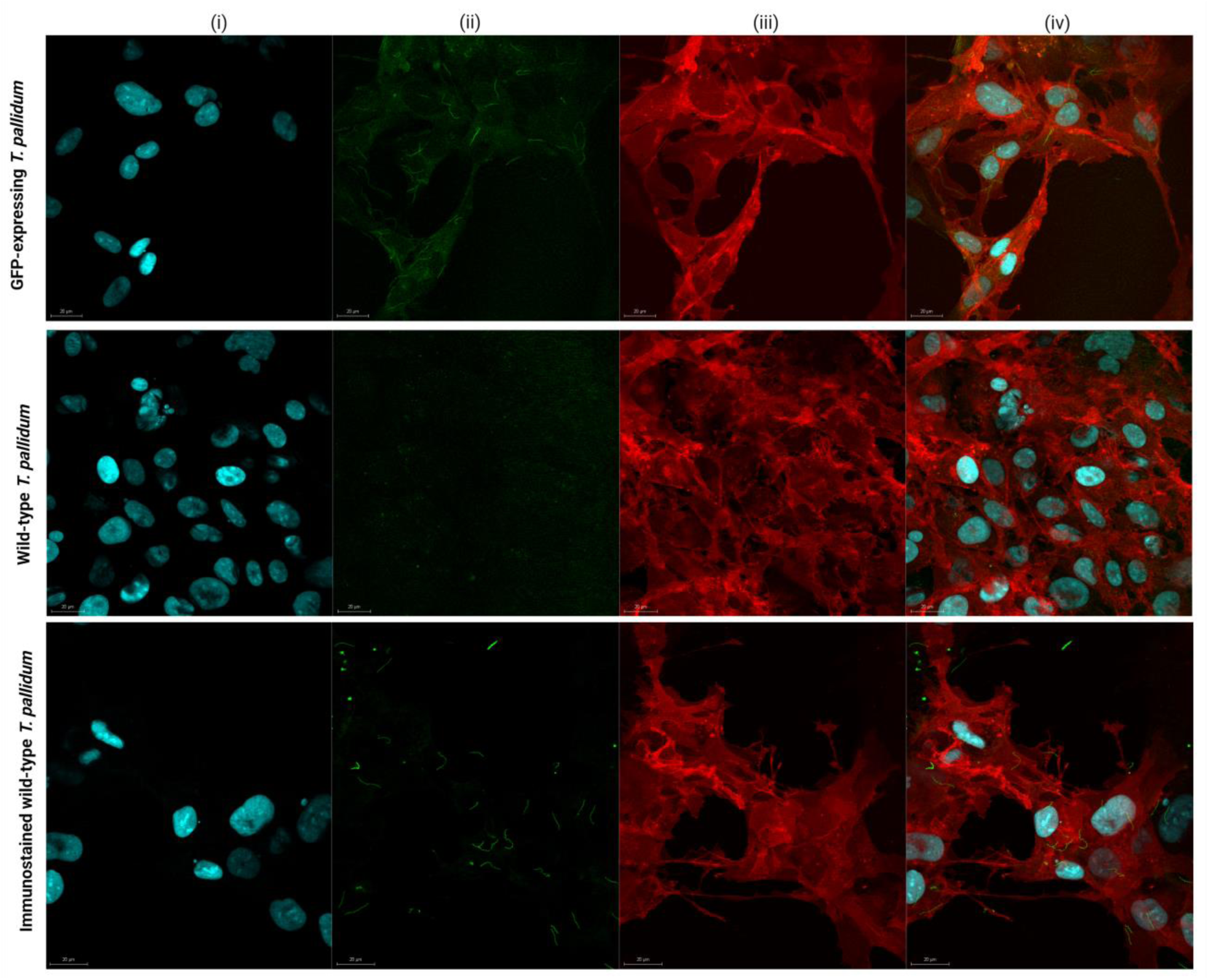
Images of GFP expressing *T. pallidum* (top panels) and wild-type *T. pallidum* (middle and bottom panels) grown on mCherry and BFP-expressing Sf1Ep cells. (i) Blue channel showing Blue Fluorescent Protein (BFP) expressed in the nuclei of Sf1Ep cells. (ii) Green channel showing Green Fluorescein Protein (GFP) expressed in GFP-SS14 *T. pallidum* cells (top panel) but not in wild type *T. pallidum* (middle panel). Bottom (ii) panel: wild-type *T. pallidum* cells visualized following FITC-immunostaining. (iii) Red channel, mCherry expressed in the membrane of Sf1Ep cells. (iv) Overlay. Scale bars: 20 μm.

### Rabbit infection

Rabbit infection was performed to further passage the GFP-SS14 strain to show evidence this strain would grow *in vivo*, and to prepare frozen stocks for further studies and dissemination. The infected rabbit developed orchitis of the left testicle on day 14 post-inoculation. In total, the GFP-SS14 treponemal yield from the animal was 1.5x10^8^ *T. pallidum* cells. A second rabbit yielded a total of 4.9x10^7^ treponemal cells after a longer time to orchitis (29 days).

## DISCUSSION

Researchers have just begun to develop molecular tools for *T. pallidum* thanks to the recent demonstration that this pathogen can internalize foreign DNA and use it as substrate for homologous recombination (3). Such evidence was used, for example, to derive the loss-of-function strain DC^KO^-SS14, which appeared to be attenuated due to its artificially impaired ability for antigenic variation (10). For the first time we have shown how *T. pallidum* can be manipulated with a *gfp-kan*^R^ cassette fusion integrated into its chromosome and express a functional GFP protein, allowing imaging of the spirochetes under a fluorescent microscope negating the need for staining with fluorophore-labeled antibodies. Fluorescence emitted by GFP-SS14 cells could also be measured using a plate reader, even though, due to the elevated number of GFP-SS14 treponemes needed to produce a signal significantly above background, and the need to eliminate Sf1Ep cells and the TpCM2 culture media, might not make direct fluorescence measurement using a plate reader a practical approach.

Eventually, syphilis investigators will benefit from the same genetic tools for visualization that helped Lyme disease research. *Borrelia* cells expressing a fluorescent tag, for example, were used to monitor expression of specific genes at different stages of the infectious cycle or in various conditions of temperature and pH *in vitro* by positioning the *gfp* gene downstream of specific promoters (12, 13). In the case of *T. pallidum*, the same approach could help identify stage- or tissue-specific differential expression of genes coding for virulence factors and putative OMPs. In the transgenic strain derived here, *gfp* is transcribed from the minimal promoter derived from the *tp0574* gene (11), which encodes a periplasmic lipoprotein involved in cell wall and envelope biogenesis (14, 15). Given its abundance, the *tp0574* message has been many times in our studies as a normalizer for gene expression (16–18). Like all *T. pallidum* genes, it remains unclear whether expression of *tp0574* is constitutive or modulated, particularly when the disease enters its latency phase. Assuming, this modulation is effected through the *tp0574* promoter, the strain derived here could help answer this question in future studies. In our case, however, studies of gene expression using a promoter-GFP reporter would be slightly complicated by the fact that no shuttle vector is yet available for *T. pallidum*, and a transgene still needs to be integrated in the pathogen’s genome. However, the presence of at least one genomic locus (*tp0009*) that does not appear to be necessary for *T. pallidum* viability or pathogenicity (3) provides the ideal target for insertions or replacements.

Another application for the GFP-SS14 strain could be its use in opsonophagocytosis (OP) experiments, performed to evaluate whether an antiserum to a putative surface-exposed antigen facilitates *T. pallidum* ingestion by activated macrophages. Following internalization, phagocytic vacuoles are generally stained with polyclonal antisera to *T. pallidum* collected from long-term infected rabbits, suggesting that *T. pallidum* antigens are not completely degraded and can be recognized within the vacuoles. The possibility that GFP expression could bypass immunostaining in OP experiments is currently being explored.

Additional possibilities include the use of the actual syphilis agent instead of a surrogate system (e.g., *Borrelia* expressing both GFP and a *T. pallidum*-specific gene) to study pathogen dissemination and adhesion, similar to Kao *et al.* (19) using live cell imaging and a Tp0751-expressing gain-of-function *Borrelia* strain to study attachment to blood vessel surfaces under fluid shear stress mimicking conditions in postcapillary venules. The live cell incubator used for these experiments, however, will need to be equipped with an appropriate containment unit and a gas exchange system to ensure maintenance of the microaerophilic environment necessary for treponemal survival throughout the experiment. For these studies, our GFP-expressing strain could be further engineered to ablate specific genes potentially involved in adhesion and tissue invasion. In addition to *in vitro* imaging, *in vivo* visualization is also being explored to study dissemination and the possibility of congenital transmission in the mouse model of infection.

Lastly, these treponemes could be used for correlative cryo-fluorescence and cryo-scanning electron microscopy as described by Strnad *et al*. (20, 21) for *Borrelia.* This imaging technique promises to enable the identification and targeting of fluorescently tagged cells with subsequent imaging at a near-to-nanometer resolution, close to their natural state, and devoid of artifacts possibly caused by chemical fixatives.

In addition, the mCherry and BFP-expressing Sf1Ep cells facilitate the visualization of the GFP-expressing *T. pallidum* and can be used to study *T. pallidum* attachment dynamics in real-time in the current *in vitro* co-culture model. Moreover, single-cell omics using isolated distinct Sf1Ep cell populations (using laser capture microdissection, for instance) can potentially uncover in detail the host-pathogen interaction and *T. pallidum* dependencies.

The *T. pallidum* strain expressing GFP opens numerous possibilities, including its use for real-time imaging of syphilis pathogens without the need for immunostaining, and for detailed studies on pathogen dissemination and host-pathogen interactions. Furthermore, the application of these tools extends beyond basic research, offering the opportunity to gain insights into gene expression, virulence factors, and the mechanics of infection, mirroring the transformative impact seen in Lyme disease research.

## EXPERIMENTAL PROCEDURES

### Ethics statement

Specific pathogen-free (SPF; *Pasteurella multocida*, and *Treponema paraluiscuniculi*) male New Zealand White (NZW) rabbits (3.5-4.5 kg in weight) were purchased from Western Oregon Rabbit Company (Philomath, OR) and housed at the University of Washington (UW) Harborview Medical Center Research and Training Building vivarium. Care was provided in accordance to the Guide for the Care and Use of Laboratory Animals (22) guidelines under a protocol approved by the UW Institutional Animal Care and Use Committee (IACUC; Protocol # 4243-01, PI: Lorenzo Giacani). Upon arrival, all animals were bled and tested with the Treponema Pallidum Particle Agglutination (TPPA; Fujirebio, Tokyo, Japan) and the Venereal Disease Research Laboratory (VDRL; Becton Dickinson, Franklin Lakes, NJ) tests to confirm lack of humoral immunity to *T. pallidum* antigens, as the vendor tests animals randomly. Both tests were performed according to the manufacturer’s instructions. Only seronegative rabbits were used for experimental infection with the GFP-SS14 strain to confirm the feasibility of *in vivo* propagation and attain stocks for storage and dissemination.

### Plasmid construct for treponemal transformation

The pUC57 vector (2,710 bp; Genscript, Piscataway, NJ) was engineered to carry a hybrid ORF composed of the *gfp* gene from *Renilla reniformis* lacking a stop codon and terminally fused to the *kan*^R^ gene from the Rts1 plasmid of *Proteus vulgaris*. The *gfp-kan*^R^ chimeric gene was positioned downstream of the *T. pallidum tp0574* gene promoter and ribosomal binding site (RBS) previously characterized by Weigel *et al.* (15). An 8-nucleotide spacer was added between the RBS and the start codon of the reporter/resistance gene in the construct. Upstream and downstream of the transgene, respectively, two homology arms corresponding to the regions flanking the *tprA* pseudogene were also cloned. The upstream arm was 998 bp-long and corresponded to genomic position 7,343-8,340 of the SS14 strain (NC021508.1/CP004011.1), which was used to derive the GFP-SS14 strain. The downstream arm was 999 bp-long and encompassed position 10,165-11,163 of the strain genome. The construct was cloned between the XheI and BamHI sites of the pUC57 vector (File S1), in opposite orientation compared to the *lac* promoter upstream of the polylinker. The insert in the final construct was Sanger-sequenced by the vendor to confirm sequence accuracy. Both the insert sequence and vector map are provided in File S1. The plasmid was transformed into TOP10 *E. coli* cells (Thermo Fisher Scientific, Waltham, MA), which were then grown in 500 ml of LB media supplemented with 100 µg/ml of ampicillin at 37⁰C inoculated from a starter culture. The plasmid was purified using the Endo-Free Plasmid Mega Kit (Qiagen, Germantown, MD) per manufacturer’s instructions. Following purification, the plasmid concentration was assessed using an ND-1000 spectrophotometer (Nanodrop Technologies, Wilmington, NC). The plasmid was then aliquoted and stored at -80⁰C until use to transform *T. pallidum*.

### Source of *T. pallidum* for *in vitro* cultivation, transformation, and selection

The SS14 strain of *T. pallidum* used to generate the GFP-expressing strain is routinely propagated in the Giacani lab according to Edmondson *et al.* (2) in the wells of a 6-well culture plate (Corning Inc, Corning, NY). This strain was isolated in 1977 in Atlanta (USA) from a secondary syphilis patient who claimed to be penicillin-allergic and who did not respond to therapy with macrolides due to the genetic resistance of the strain to this class of antibiotics, which was however unknown at that time. This strain was originally provided to Dr. Sheila Lukehart at the University of Washington by Dr. Sandra A. Larsen, Centers for Disease Control and Prevention, Atlanta, GA. The SS14 strain is used in research settings in addition to the Nichols strain as the laboratory isolate for the SS14 clade of circulating *T. pallidum* strains (23).

Transformation of *T. pallidum* was performed as previously reported (24). Briefly, to prepare the treponemal inoculum for transformation, one week-old co-cultures of Sf1Ep cells and treponemes were trypsinized and spirochetes were enumerated via dark-field microscopy (DFM). A total of 7x10^7^ treponemes were inoculated on wells of a 24-well plate containing 2.5x10^4^ Sf1Ep cells seeded the previous day. Following treponemal addition, the total volume of media in each well was brought to 2.5 ml. Two days following the addition of treponemes a full media change was performed. Two days after that, the culture media was removed gently so as not to disturb Sf1Ep cells and adherent treponemes.

Media was then replaced with 500 µl of transformation buffer containing 10 µg total of plasmid carrying the transgene. Cells were incubated in transformation buffer for 10 min at 34°C in the microaerophilic atmosphere (MA) incubator and then washed twice with equilibrated TpCM-2 media to remove free plasmid. Finally, 2.5 ml of fresh TpCM-2 equilibrated in MA were added to the wells, and plates were returned to the MA incubator. The following day, kanamycin sulfate (Sigma-Aldrich, St. Louis, MO) was added to a final concentration of 200 µg/ml. As controls, wild-type treponemes processed as above but not transformed were incubated in fresh TpCM-2 media containing 200 µg/ml of kanamycin sulfate and, in parallel, wild-type treponemes were propagated in antibiotic-free media as a control. Kanamycin sulfate-containing TpCM-2 media was exchanged weekly but treponemes were sub-cultured once every two weeks to minimize loss of cells until they reached a density of ∼10^7^ cells/ml, per DFM counting. At this point, the culture was expanded as per protocol (2) into all the wells of a 6-well plate to further expand the strain prior to whole genome sequencing and rabbit inoculation.

### Fluorescence measurement

For GFP fluorescence measurements, WT and GFP-expressing *T. pallidum* cells were harvested from plates and counted using DFM. Cells were pelleted by centrifugation for 10 minutes at 20,000 x g on a tabletop centrifuge. Supernatants were removed, and cells resuspended in PBS. 300 µl of each suspension was transferred to the wells of a black OptiPlate-96F (Revvity, Waltham, MA) for top fluorescence reading. Excitation and emission wavelength were 485 and 528 nm, respectively, with a gain of 120 and readings were performed in a Synergy HTX multi-mode plate reader with the 3.10.6 version of the Gen5 Microplate Reader and Imager Software (BioTek, Winooski, VT). Reported data represent fluorescence readings (expressed in Relative Fluorescence Units, RFU) ± SEM of triplicate experimental wells. Resulting values were graphed using Prism software (Version 9.5.1; GraphPad, San Diego, CA). Differences between fluorescence level were compared using student’s t-test, with significance set at p<0.05.

### DNA extraction and whole-genome sequencing

DNA extraction from cultured treponemes propagated in 6-well plates following transformation and expansion was performed using the QIAamp DNA mini kit (Qiagen) according to the manufacturer’s instructions. Extracted DNA was stored at -80°C until use for qualitative PCR to assess the transgene integration in the *tprA* locus or whole genome sequencing (WGS). Primers and amplicon size are reported in Table 1. These amplifications used primers targeting, respectively, the *tprA* locus (deleted in the GFP-expressing strain), the *kan*^R^ cassette (absent in the wild-type strain), and the *tp0574* gene (present in both strains). An additional amplification targeted the region of the *T. pallidum* SS14 genome immediately upstream and downstream of the *tprA* homology arm of the vector (and hence not cloned into the transformation vector) with the rationale that, if successfully integrated in lieu of the *tprA* pseudogene, the transgene would generate an amplicon with a restriction site for the XhoI enzyme, not present in the wild-type strain. Amplification conditions used were previously reported (24). Restriction digestion with the XhoI (New England Biolabs, Ipswich, MA) was performed according to the manufacturer’s instruction.

For whole genome sequencing (WGS), pre-capture libraries were prepared from up to 100 ng of input DNA using the KAPA Hyperplus kit (Roche) and TruSeq adapters and barcoded primers (Illumina), following the manufacturer’s protocols, yielding an average fragment size longer than 500 bp. Hybrid capture of *T. pallidum* genomic DNA was performed overnight (>16 hours) using a custom IDT xGen panel designed against the reference genome NC_ 010741, following the manufacturer’s protocol. Libraries were sequenced on a Nextseq 2000 (Illumina). Paired end 2x151 bp reads were adapter and quality trimmed using Trimmomatic v0.39 (25), and mapped using Bowtie2 (26) to the wild type SS14 reference (NC_021508.1) and a custom GFP-SS14 genome reference in which the *tprA* pseudogene (TPASS_RS00040; *tp0009*) was replaced with the *gfp-kan*^R^ reporter/resistance transgene preceded by the *tp0574* promoter. Manual confirmation of expected coverage and junctions was performed by visual inspection in Geneious Prime v2020.1.2 (27). Consensus sequences were generated as previously described (9). SS14-GFP and SS14-WT reads, and consensus sequences are available in the NCBI SRA in BioProject PRJNA1088509.

### Plasmid construct for Sf1Ep cell transduction

Recombinant lentivirus with the custom vector (ID: VB230919-1538krx) was ordered from VectorBuilder Inc. (Chicago, IL). Detailed sequence information and vector map are available at https://en.vectorbuilder.com/design/retrieve.html using the above vector ID. For this construct, we selected the EF1A promoter with MLS-mCherry and NLS-BFP2, separated by a T2A peptide for co-translational cleavage of the encoded polypeptide. MLS-mCherry contains a signal peptide that targets mCherry to the membrane, and NLS-BFP2 contains a signal peptide that directs the BFP to the nucleus. A puromycin resistance gene was added as a selection marker, linked with the fluorescence markers via IRES to ensure that only fluorescent cells are selected. mCherry was intentionally chosen to select for the most distinct fluorophore compared to GFP. This system can be applied to other mammalian cell lines, and the colors can be targeted to different cellular compartments.

### Transduction and selection of mCherry and BFP-expressing Sf1Ep cells

Sf1Ep cells (CRL-6502 ™, passage 21) were seeded in a 6-well plate one day prior to transduction in the concentration of 10^5^ cells/well. The following day, when the cells reached 30%-50% confluence, the old media was removed, and 1 ml of the viral suspension corresponding to an MOI of 10 was added. After 24 hours, the media was replaced, and once confluent (after 7 days), one million cells were transferred to a well of the 6-well plate containing 3 μg/ml of puromycin. After a week, the cells were transferred to a T75 flask and treated with 3 µg/ml of puromycin until selection was complete.

### Sample preparation and fluorescence microscopy

One hundred thousand mCherry and BFP-expressing Sf1Ep cells were seeded on the sterile circular coverslip (12 mm, Paul Marienfeld GmbH & Co.KG, Lauda-Königshofen, Germany) placed in the 24-well plate (one slide per well) in the media and cultured overnight in the incubator. After 24 hours, the cells were washed, and TPCM-2 was added, and the cells were equilibrated in a low-oxygen incubator. After 3 hours, 10^6^ GFP-expressing *T. pallidum* cells and 10^6^ WT cells were added to the cells grown on coverslips and left in the low oxygen incubator for 24 hours. Then, the media was removed, and the samples were fixed with 4% paraformaldehyde (PFA, Thermo Fisher Scientific) for 60 minutes. Afterwards, PFA was removed, and the samples were washed with PBS. For the immunostaining of wild-type *T. pallidum* cells, we incubated the coverslips with 10% bovine serum albumin solution (BSA, Merck Life Science UK Limited, Gillingham, UK) for 30 minutes to block non-specific interactions, followed by 1 hour of incubation with FITC anti-*Treponema pallidum* antibody (abcam, Cambridge, UK) diluted 1:100 in PBS with 1% BSA and 2% Triton™ X-100 (Merck Life Science UK Limited, Gillingham, UK). The incubation was performed in the dark with shaking. After incubation, the cells were washed twice with 1% BSA. All coverslips were then mounted onto microscope slides with a drop of diamond anti-fade reagent (ProLong Diamond Antifade Mountant; Thermo Fisher Scientific) and kept in the dark overnight. Images were acquired on Leica TCS SP8 microscope (Leica Microsystems, Milton Keynes, UK) using 63x oil objective. LAS X Life Science Microscope Software Platform was used to export the images.

### Rabbit infection

Intratesticular infection of naïve rabbits was performed as previously described (28). Briefly, one rabbit was injected with 10^7^ treponemes/testis and a second one with a slightly lower dose (4.6x10^6^/testis). Both rabbits were subsequently monitored twice weekly to assess development of orchitis via palpation of the testes. Prior to euthanasia, the presence of treponemes was assessed in fluid from a testicular needle aspirate using DFM.

## ACKNOWLEDGEMENTS

This work was supported by the Open Philanthropy Pledge #8394150 (to Lorenzo Giacani, University of Washington), by the National Institute for Allergy and Infectious Diseases (NIAID) grant number U19AI144133 (Genomics and Isolation Core; Core leaders Alexander L. Greninger and Lorenzo Giacani. PI: Anna Wald, University of Washington), and by the Biotechnology and Biological Sciences Research Council (BBSRC) Discovery Fellowship (grant number BB/010589) (to Linda Grillová) and Wellcome grant #206194 (to Nick Thomson, George Lacey and David Goulding, Wellcome Sanger Institute). The content of this study is solely the responsibility of the authors and does not necessarily represent the official views of Open Philanthropy, NIAID, or the BBSRC. The funders had no role in study design, data collection, and analysis, decision to publish, or preparation of the manuscript.

## AUTHOR CONTRIBUTIONS

L.G. (University of Washington): Conceived the experimental design for *T. pallidum* transformation, designed the transgene, oversaw all experiments at the University of Washington, wrote the manuscript. E.R.: performed the *T. pallidum* transformation experiment; L.T.: propagated treponemes; A.T.T. performed molecular assays. B.J.M. and L.H.X.: performed rabbit work. N.A.P.L. and A.L.G.: performed genome sequencing and genomic data analysis. All authors reviewed and provided comments on the manuscript prior to submission. L.G. (Wellcome Sanger Institute): conceived the experimental design for Sf1Ep cells transduction, performed and optimized cell selection and sample preparation for the microscopy, oversaw all experiments at the Wellcome Sanger Institute, wrote the manuscript. G.L.: propagated treponemes and performed fluorescent microscopy. D.G.: performed fluorescent microscopy. N.T.: supervised the project at the Wellcome Sanger Institute. All authors reviewed and provided comments on the manuscript prior to submission.

## Notes

### Competing Interest Statement

The authors have declared no competing interest.

## REFERENCES

1. Nichols HJ, Hough WH. Demonstration of *Spirochaeta pallida* in the cerebrospinal fluid. JAMA. 1913;60:108–10.

2. Edmondson DG, Hu B, Norris SJ. Long-Term In Vitro Culture of the Syphilis Spirochete *Treponema pallidum* subsp. *pallidum*. mBio. 2018;9(3).

3. Romeis E, Tantalo L, Lieberman N, Phung Q, Greninger A, Giacani L. Genetic engineering of *Treponema pallidum* subsp. *pallidum*, the Syphilis Spirochete. PLoS Pathog. 2021;17(7):e1009612.

4. Haynes AM, Giacani L, Mayans MV, Ubals M, Nieto C, Pérez-Mañá C, et al. Efficacy of linezolid on *Treponema pallidum*, the syphilis agent: A preclinical study. EBioMedicine. 2021;65:103281.

5. Tantalo LC, Lieberman NAP, Pérez-Mañá C, Suñer C, Vall Mayans M, Ubals M, et al. Antimicrobial susceptibility of *Treponema pallidum* subspecies *pallidum*: an in-vitro study. Lancet Microbe. 2023;4(12):e994–e1004.

6. Edmondson DG, Wormser GP, Norris SJ. In Vitro Susceptibility of Treponema pallidum subsp. pallidum to Doxycycline. Antimicrob Agents Chemother. 2020;64(10).

7. Hayes KA, Dressler JM, Norris SJ, Edmondson DG, Jutras BL. A large screen identifies beta-lactam antibiotics which can be repurposed to target the syphilis agent. npj Antimicrobials and Resistance. 2023;1(1):4.

8. De Lay BD, Cameron TA, De Lay NR, Norris SJ, Edmondson DG. Comparison of transcriptional profiles of *Treponema pallidum* during experimental infection of rabbits and in vitro culture: Highly similar, yet different. PLoS Pathog. 2021;17(9):e1009949.

9. Lin MJ, Haynes AM, Addetia A, Lieberman NAP, Phung Q, Xie H, et al. Longitudinal TprK profiling of in vivo and in vitro-propagated *Treponema pallidum* subsp. *pallidum* reveals accumulation of antigenic variants in absence of immune pressure. PLoS Negl Trop Dis. 2021;15(9):e0009753.

10. Romeis E, Lieberman NAP, Molini B, Tantalo LC, Chung B, Phung Q, et al. *Treponema pallidum* subsp. *pallidum* with an Artificially impaired TprK antigenic variation system is attenuated in the Rabbit model of syphilis. PLoS Pathog. 2023;19(3):e1011259.

11. Weigel LM, Brandt ME, Norgard MV. Analysis of the N-terminal region of the 47-kilodalton integral membrane lipoprotein of *Treponema pallidum*. Infect Immun. 1992;60(4):1568–76.

12. Carroll JA, Stewart PE, Rosa P, Elias AF, Garon CF. An enhanced GFP reporter system to monitor gene expression in *Borrelia burgdorferi*. Microbiology (Reading). 2003;149(Pt 7):1819–28.

13. Gautam A, Hathaway M, McClain N, Ramesh G, Ramamoorthy R. Analysis of the determinants of bba64 (P35) gene expression in *Borrelia burgdorferi* using a gfp reporter. Microbiology (Reading). 2008;154(Pt 1):275–85.

14. Deka RK, Machius M, Norgard MV, Tomchick DR. Crystal structure of the 47-kDa lipoprotein of *Treponema pallidum* reveals a novel penicillin-binding protein. J Biol Chem. 2002;277(44):41857–64.

15. Weigel LM, Radolf JD, Norgard MV. The 47-kDa major lipoprotein immunogen of *Treponema pallidum* is a penicillin-binding protein with carboxypeptidase activity. Proc Natl Acad Sci U S A. 1994;91(24):11611–5.

16. Haynes AM, Fernandez M, Romeis E, Mitjà O, Konda KA, Vargas SK, et al. Transcriptional and immunological analysis of the putative outer membrane protein and vaccine candidate TprL of Treponema pallidum. PLoS Negl Trop Dis. 2021;15(1):e0008812.

17. Giacani L, Brandt SL, Ke W, Reid TB, Molini BJ, Iverson-Cabral S, et al. Transcription of TP0126, *Treponema pallidum* putative OmpW homolog, is regulated by the length of a homopolymeric guanosine repeat. Infect Immun. 2015;83(6):2275-89.

18. Giacani L, Molini B, Godornes C, Barrett L, Van Voorhis WC, Centurion-Lara A, et al. Quantitative analysis of *tpr* gene expression in *Treponema pallidum* isolates: differences among isolates and correlation with T-cell responsiveness in experimental syphilis. Infect Immun. 2007;75(1):104–12.

19. Kao WA, Pětrošová H, Ebady R, Lithgow KV, Rojas P, Zhang Y, et al. Identification of Tp0751 (Pallilysin) as a *Treponema pallidum* Vascular Adhesin by Heterologous Expression in the Lyme disease Spirochete. Sci Rep. 2017;7(1):1538.

20. Strnad M, Elsterová J, Schrenková J, Vancová M, Rego RO, Grubhoffer L, et al. Correlative cryo-fluorescence and cryo-scanning electron microscopy as a straightforward tool to study host-pathogen interactions. Sci Rep. 2015;5:18029.

21. Dahlberg PD, Moerner WE. Cryogenic Super-Resolution Fluorescence and Electron Microscopy Correlated at the Nanoscale. Annu Rev Phys Chem. 2021;72:253–78.

22. Guide for the Care and Use of Laboratory Animals: The National Academic Press; 2011.

23. Lieberman NAP, Lin MJ, Xie H, Shrestha L, Nguyen T, Huang ML, et al. *Treponema pallidum* genome sequencing from six continents reveals variability in vaccine candidate genes and dominance of Nichols clade strains in Madagascar. PLoS Negl Trop Dis. 2021;15(12):e0010063.

24. Phan A, Romeis E, Tantalo L, Giacani L. In Vitro Transformation and Selection of *Treponema pallidum* subsp. *pallidum*. Curr Protoc. 2022;2(8):e507.

25. Bolger AM, Lohse M, Usadel B. Trimmomatic: a flexible trimmer for Illumina sequence data. Bioinformatics. 2014;30(15):2114–20.

26. Langmead B, Salzberg SL. Fast gapped-read alignment with Bowtie 2. Nat Methods. 2012;9(4):357–9.

27. Kearse M, Moir R, Wilson A, Stones-Havas S, Cheung M, Sturrock S, et al. Geneious Basic: an integrated and extendable desktop software platform for the organization and analysis of sequence data. Bioinformatics. 2012;28(12):1647–9.

28. Lukehart SA, Marra CM. Isolation and laboratory maintenance of *Treponema pallidum*. Curr Protoc Microbiol. 2007;Chapter 12:7:12A.1.1–A.1.8.

