## Supplementary material for "Bright New Resources for Syphilis Research: Genetically Encoded Fluorescent Tags for *Treponema pallidum* and Sf1Ep Cells": File S1.docx

**gfp-*kan*^R^ plasmid insert**

*tprA* upstream flanking region/homology arm

*Tp0574* promoter/ribosomal binding site

gfp gene

*kan*^R^ gene

*tprA* downstream flanking region/homology arm

EcoRI (top, GAATTC) and Xbal (bottom, TCTAGA) restriction sites for cloning into pUC57 vector

XhoI site (CTGGAG) within the Kan^R^ cassette

GAATTCTATAAAGGCCATTGTGGTTAGTCAGGCAGTTCCCGGCGTTTCAAAGGCGTTTGGGATCATTAAGTCTAAACGTCCTGATGTTTTGCTTTTTGCGGGAGAACCACTTGAGCCGGTAGAGATGCTGCAGGAGTCTGCAGACATCGTGGTCAGTCAGGACTACTTGTTCGGTGGATATGCCGTTCCGTGGGTTGCGGAAAGGATGGGGGCGCGCACATTGGTGCATGTCTCTTTTCCCCGGCATATGTCCTACCCCGGTTTGAGGGTTAGGCGTACGGTGATGAGGGCAGCATGTACCGATTTGGGACTTTCCTTCGCACACGAGGAAGCGCCTGATCCTGTAGACGGTGTCAGTGACGGAGAACTTGAGGATTTTTTCCACAAGACGATTGTGAAGTGGATCAAAAAATATGGCAAGGAAACCCTGTTCTACTGCACCAATGACGCTCACAACAGGCCGCTCATCAGTGCCTTGTTGAAATATGGCGGTATGCTAATTGGTGCAACCATCTTCGATTACGCTGATGCGCTCGGGGTGCATTATGCTGAGCTTGAAGACGTGTATAAAATACGAGAGAAGGTTGAGAAGTCATTGGTTGCCTTCGGCGCAGAGGGGCGCTTTGGATTAAATTTAAATGCACAGGCATTTACGGTGACCATGGGTTTTGTGGAGTATGCGCGCAAAATCATAGATGGCGAACCGCGTAAAGATGATATGCGTGAAGCTCTTGCCGAATCCTTCGACTTGTTTACGCGTGACGCACATTGGCGTATTGCTCCTTACCTAAGACTGAAAACGCACGAAATTGTTCCGAATCACGTGCTGGTGTATACGGACACATACGTCCTGGGTAAATTTACCTTGCCCGTCACAGACCAAGTACTCCCAGAAGGGTATTGGGCATTGACCGCTAAGGAATAAGAACTCCGTTCGGGTTTTCTGTTTGTAGCCGGGGAGATGGATCGCTTTCTCTGTTTGGCAATGTCGCCGTCTCCCTGGGAGCGGATCCTCCCAAAAAGAGGAAGGACGCGCCTGTGTGTGCTCTGCATAAGACGTTGACAATCCCTGTGGGGCGTGCCTATACTCAGGCCCTCTATACGGAGGTGTAATCATGAGTAAAGGAGAAGCACTTTTCACTGGAGTTGTCCCAATTCTTGTTGAATTAGATGGTGATGTTAATGGGCACAAATTTTCTGTCAGTGGAGAGGGTGAAGGTGATGCAACATACGGAAAACTTACCCTTAAATTTATTTGCACTACTGGAAAACTACCTGTTCCATGGCCAACACTTGTCACTACTCTTACGTATGGTGTTCAATGCTTTTCAAGATACCCAGATCATATGAAACGGCATGACTTTTTCAAGAGTGCCATGCCCGAAGGTTATGTACAGGAAAGAACTATATTTTTCAAAGATGACGGGAACTACAAGACACGTGCTGAAGTCAAGTTTGAAGGTGATACCCTTGTTAATAGAATCGAGTTAAAAGGTATTGATTTTAAAGAAGATGGAAACATTCTTGGACACAAATTGGAATACAACTATAACTCACACAATGTATACATCATGGCAGACAAACAAAAGAATGGAATCAAAGTTAACTTCAAAATTAGACACAACATTGAAGATGGAAGCGTTCAACTAGCAGACCATTATCAACAAAATACTCCAATTGGCGATGGCCCTGTCCTTTTACCAGACAACCATTACCTGTCCACACAATCTGCCCTTTCGAAAGATCCCAACGAAAAGAGAGACCACATGGTCCTTCTTGAGTTTGTAACAGCTGCTGGGATTACACATGGCATGGATGAACTATACAAGTCCGGACTCATGAGCCATATTCAACGGGAGACGTCTTGCTCGAGGCCGCGATTAAATTCCAACCTGGATGCTGATTTATATGGGTATAGATGGGCTCGCGATAATGTCGGGCAATCAGGTGCGACAATCTATCGATTGTATGGGAAGCCCGATGCGCCAGAGTTGTTTCTGAAACATGGCAAAGGTAGCGTTGCCAATGATGTTACAGATGAGATGGTCAGACTAAACTGGCTGACGGCATTTATGCCTCTTCCGACCATCAAGCATTTTATCCGTACTCCTGATGATGCATGGTTACTCACCACTGCGATCCCCGGGAAAACAGCATTCCAGGTATTAGAAGAATATCCTGATTCAGGTGAAAATATTGTTGATGCGCTGGCAGCGTTCCTGCGCCGGTTGCATTCGATTCCTGTTTGTAATTGTCCTTTTAACAGCGATCGCGTATTTCGTCTCACTCAGGCGCAATCACGAATGAATAACGGTTTGGTTGATGCGAGTGATTTTGATGACGAGCGTAATGGCTGGCCTGTTGAACAAGTCTGGAAAGAAATGCATAAGCTTTTGCCATTCTCACCGGATTCAGTCGTCACTCATGGTGATTTCTCACTTGATAACCTTATTTTTGACGAGGGGAAATTAATAGGTTGTATTGATGTTGGACGAGTCGGAATCGCAGACCGATACCAGGATCTTGCCATCCTATGGAACTGCCTCGGTGAATTTTCACCTTCATTACAGAAACGGTTTTTTTATAAATATGGCATTGATAATCCTGATATGAATAAATTGCAGTTTCATTTGATGCTCGATGAGTTTTTCTGAAGTTTAGTACAACGATGTCATGTGTCAGATCTAGCAGTATCTGTAATGTATGTTGGTGTACATTAGATATTCGTGGGTGGGAAGAAGAGTCACTTTCTGGGGAGGCGTATAGAAGGAACGGGGCGTGGTGTTGTGCCATTTGTGCGAAAACTGAGTGAAGTAGTGAAAAAAATTACCGCTGACGGGAAAAAATGTTGATCGTGTTTATGAAAGGGTCATAATGGCTGCCCTATGGGCGCCTGTATATCCGTATATGCGCGTTTTGCGTTAGGGTGTGGGGTGTTTTTCCTTCATGGTGCGGTTTTGGACGGGGTTTCACGCGCCTTTTCGTCCTCCGCCGCGTTCAGCGGTTCTGCTGAACTTAGCTGGGGTGTCGTCTTTGATGCAGAAGGTGCCTCTCCAGTTACAGCGGGTAAAAGCATACGACATGGGTTTCGCACGAAGAGCAGCTGGAAGCTTGCTTTTCCCTTGTTGCCCAAGAAAGGCGCCACGTATACGAGCTTTTCAGGTGAGGATCCCATATGGGTTGAGCTTTCTCTCAAGGGATTGAAGGTGGATTTTGAAAGTGCTTTAGGGTCGGGAACTGCGGATCCAAGTATGACGACGCGTTCTCCTTTCTTAAAGTCAGGAAGAAGCGATTTTTCCCTTGAGGCCACACTCCACCTCTACGATGTCTCTTTTTCTGTAGGAAAAGATCCCGTTTTTCCCTCTAATTTTGCGCAGTTGTGGACCCCCTTTATTACTACTAGTTATGAGTCAAGGAGCGTCAAATACGCTCCAGGGTTTGGTGGGGTTGGCGGAAAAATCGCATATCAGGCACGGAATATTTCGAACAGTGGCATTACATTCAACTGTGCCCTTTCCTTTTCGTCGAACGGTATATGGAAAAGTGCTCCTTCTGTCACCTCTAAGGTGAAAGGAAAGGGCACCAATAGTCGGCGCATGCCAGCGGTCTAGA

**TprA-RGFP-Kan_pUC57 plasmid map**

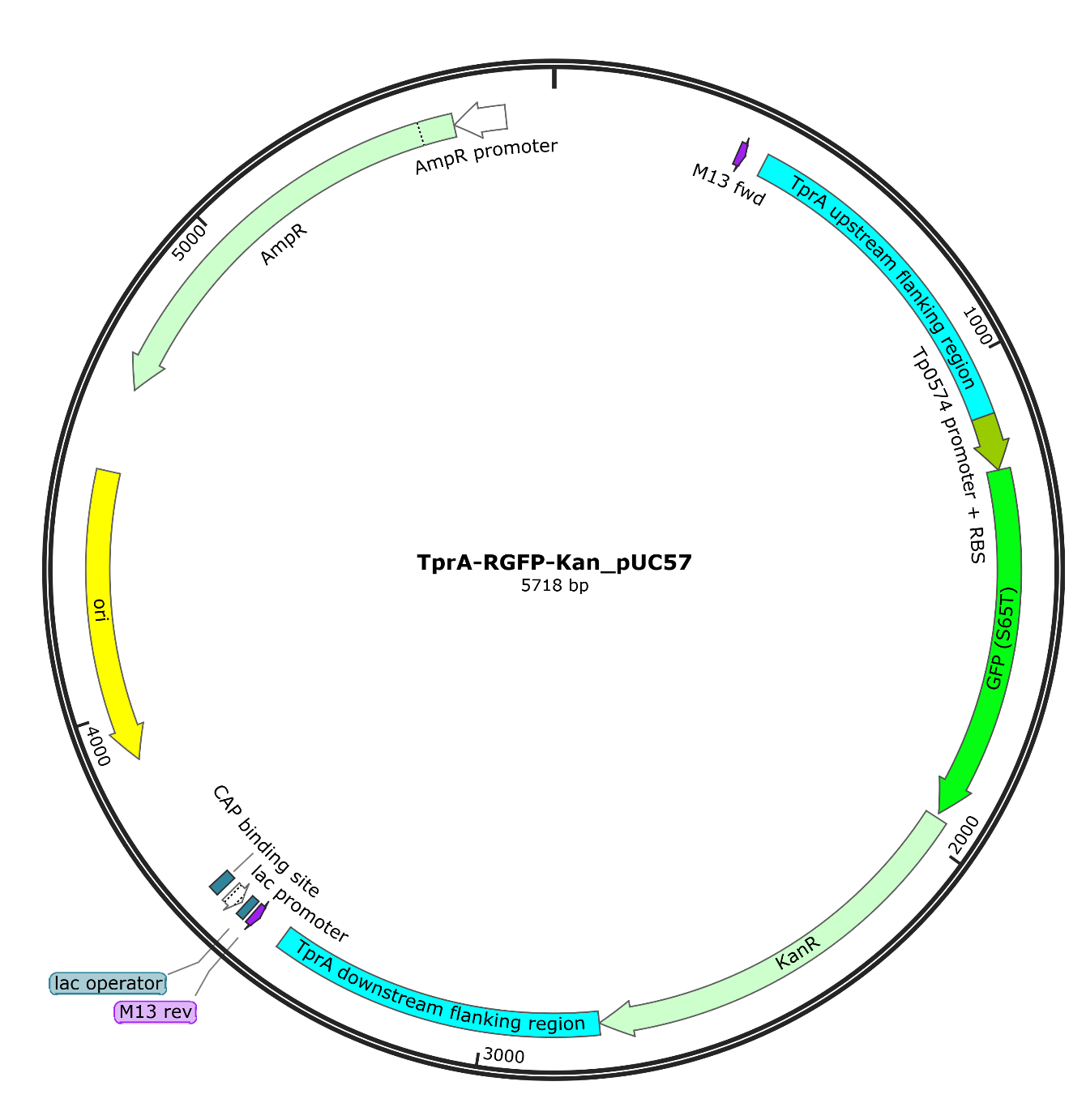

**Map of the pUC57 Vector carrying the chimeric *gfp-kan*^R^ gene.** The vector employs the tp0574 promoter/RBS to drive transcription of a hyring gfp-kanR gene positioned between two ~1Kbp homology arms guiding recombination into the *tprA* (*tp0009*) locus of *T. pallidum*.

**Recombinant lentivirus vector map**

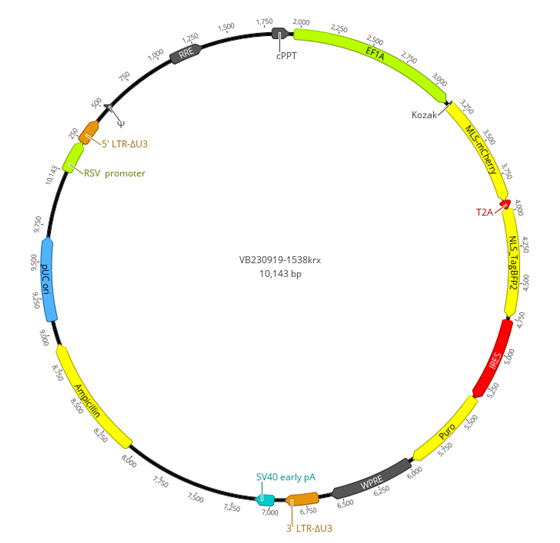

**Recombinant Lentivirus Vector Design VB230919-1538krx.** The vector employs the EF1A promoter to drive the expression of two fluorescent proteins: MLS-mCherry, targeted to the membrane of Sf1Ep cells, and NLS-BFP2, directed to the nucleus, separated by a T2A peptide for co-translational cleavage. A puromycin resistance gene, linked via IRES, serves as a selection marker. Additional genes required for lentiviral packaging were incorporated by VectorBuilder Inc.
